# Inter-scanner brain MRI volumetric biases persist even in a harmonized multi-subject study of multiple sclerosis

**DOI:** 10.1101/2022.05.05.490645

**Authors:** Kelly A. Clark, Carly M. O’Donnell, Mark A. Elliott, Shahamat Tauhid, Blake E. Dewey, Renxin Chu, Samar Khalil, Govind Nair, Pascal Sati, Anna DuVal, Nicole Pellegrini, Amit Bar-Or, Clyde Markowitz, Matthew K. Schindler, Jonathan Zurawski, Peter A. Calabresi, Daniel S. Reich, Rohit Bakshi, Russell T. Shinohara, the NAIMS Cooperative

## Abstract

**Background/Purpose:** Multicenter study designs involving a variety of MRI scanners have become increasingly common. However, these present the issue of biases in image-based measures due to scanner or site differences. To assess these biases, we imaged 11 volunteers with multiple sclerosis (MS) with scan and rescan data at 4 sites.

**Materials and Methods:** Images were acquired on Siemens or Philips scanners at 3-tesla. Automated white matter lesion detection and whole brain, gray and white matter, and thalamic volumetry were performed, as well as expert manual delineations of T1 and T2 (FLAIR) lesions. Random effect and permutation-based nonparametric modeling was performed to assess differences in estimated volumes within and across sites.

**Results:** Random effect modeling demonstrated model assumption violations for most comparisons of interest. Non-parametric modeling indicated that site explained > 50% of the variation for most estimated volumes. This expanded to > 75% when data from both Siemens and Philips scanners were included. Permutation tests revealed significant differences between average inter- and intra-scanner differences in most estimated brain volumes (*P* < .05). The automatic activation of spine coil elements during some acquisitions resulted in a shading artifact in these images. Permutation tests revealed significant differences between thalamic volume measurements from acquisitions with and without this artifact.

**Conclusion:** Differences in brain volumetry persisted across MR scanners despite protocol harmonization. These differences were not well explained by variance component modeling; however, statistical innovations for mitigating inter-scanner differences show promise in reducing biases in multi-center studies of MS.

## Introduction

Brain volumetry performed on magnetic resonance (MR) images is common practice for monitoring disease status and progression in patients with multiple sclerosis (MS). Disease course and longitudinal outcomes are often assessed by determining changes in hypointense lesions on T1-weighted images and hyperintense lesions on T2 (FLAIR) images. In addition, volumetric changes across different brain structures are used to quantify atrophy.

Previous studies have highlighted the importance of imaging biomarkers in the diagnosis, management, and therapeutic trial investigation of MS. The formation of white matter lesions in the brain is an established hallmark of MS pathogenesis, and their presentation on MR images is employed in the differential diagnosis of MS from other disease mimics.^1^ The formation and persistence of T1-hypointense (“black hole”) lesions, which suggest axonal loss and tissue destruction, have been associated with disability and disease progression in MS patients.^2–4^ Brain volume changes, notably whole brain atrophy, which are quantified in measurements obtained from MR images, have been shown to be correlated with disability progression in patients with MS.^5,6^ Rates of gray matter atrophy have been shown to differ across different stages of MS,^7^ and both white and gray matter atrophy have been linked to the development of neuropsychological symptoms and cognitive impairment in patients with MS.^8^

Accurately estimating brain volumes and lesion load is crucial for defining disease status to evaluate individual patients and assess the efficacy of therapies in clinical trials; however, volumetric measurements have been shown to vary across a subject’s measures even under ideal research conditions where scanning technique (equipment and pulse sequence parameters), and duration of follow-up are carefully controlled.^9^ Previous studies that explored site or scanner effect in either healthy people or people living with Alzheimer’s disease have reported mixed results, with some demonstrating that the use of multiple scanners has the potential to exacerbate biases in estimated average brain volumes and negatively affect the reproducibility of various measures across different sites and scanners,^10,11^ while others have demonstrated high reproducibility regardless of scanner.^12–14^

Site-related biases in brain volumes and methods for mitigating such bias including protocol standardization and image harmonization have been investigated in patients living with MS,^15–17^ yet much remains to be understood regarding the full extent of how site affects variation in images when MS pathology is present.

Multicenter studies involving a variety of MR scanners are becoming increasingly common to meet the needs of larger sample sizes and increased statistical power in research settings. Therefore, it is vital to understand expected intra- and inter-scanner differences and their impact on volumetry in people with MS to properly design clinical trials and accurately assess MRI results. Our published pilot study, which investigated scanner and methodological variation using a standardized protocol on one traveling participant with MS, revealed that greater than 50% of the variation of most brain ROI volumes could be explained by site. Additionally, differences in lesion volumes across sites were as large as 25%.^15^

To further assess variation in brain volumes attributable to site, here we imaged 11 volunteers with stable MS using a harmonized protocol developed by the North American Imaging in Multiple Sclerosis (NAIMS) Cooperative at 4 different sites. The NAIMS Cooperative was established in 2012 to unite experts in the field of MS imaging to improve MS research and routine practices through several initiatives, including the creation of standard imaging protocols and the identification of reliable imaging markers to monitor disease and therapeutic outcomes.^18,19^ Here, we explore average inter- and intra-scanner differences across volumes for several different regions of interest (ROIs) in the brain obtained using a variety of automated volumetry methods and manual lesion segmentation.

## Materials and Methods

### Participants

Eleven people with stable MS were recruited to participate in the study across all four sites. Nine of these participants were imaged at all 4 study centers, which included the University of Pennsylvania (Penn), the Brigham and Women’s Hospital (Brigham), the National Institute of Health (NIH), and the Johns Hopkins University (JHU). 2 participants were imaged at 2 and 3 sites, respectively, due to a pause in research activity during the COVID-19 pandemic. The mean age of our 11 participants (4 male, 7 female) at time of enrollment was 38 (range 29-47) years. Expanded Disability Status Scales of the 10 participants who underwent clinical assessment ranged from 0 to 3 with a mean score of 1.9 and a median score of 2.25, and the median timed 25 walk was 5.6 seconds. All participants were receiving established disease modifying treatments (DMTs) at the time of study enrollment. DMTs were not changed for at least 6 months prior to enrollment. Each participant signed an informed consent form for this study, which was approved by the University of Pennsylvania’s institutional review board (IRB). Brigham, NIH, and JHU ceded to Penn’s central IRB through a reliance agreement.

### Imaging

A standardized, high-resolution, 3-tesla (3T) MRI brain scan protocol developed by the NAIMS Cooperative was performed at each site. The protocol was compliant with recent internationally issued guidelines.^19^ Images were acquired on Siemens Skyra (Brigham, NIH), Siemens Prisma (Penn), or Philips Achieva (JHU) scanners. Scan-rescan image pairs were acquired on the same day at each visit and participants were removed from the scanner and repositioned between scans. 3D T1-weighted and 3D T2-FLAIR images acquired at each site for a single participant are shown in **Figure 1**.

**Figure 1.**
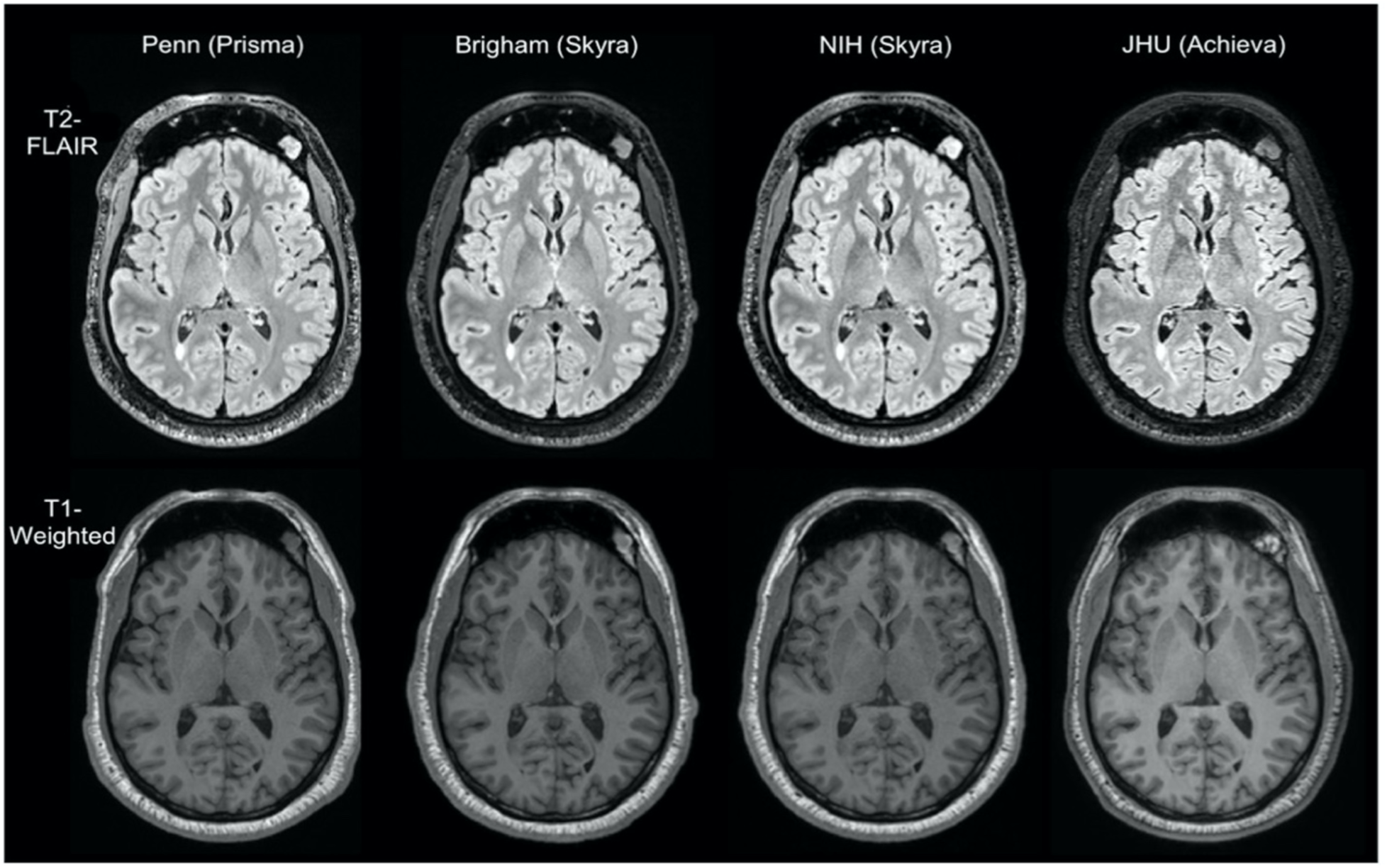
3D T1-weighted and T2-FLAIR images acquired during different sessions on different scanners for one participant.

Gradient non-linearity (GNL) has been shown to increase variation in upper cervical cord area volumes in people living with MS by as much as 15%, and GNL distortion correction methods have been shown to minimize this variation ^20^, so an additional pair of geometric distortion-corrected scan and re-scan images were reconstructed at each of the Siemens sites to compare with images without distortion correction. Relevant acquisition sequences used in this analysis are shown in the **Table 1**.

**Table 1:** 3T brain MRI anatomic acquisition protocols^a^.

| **Table1 : 3T brain MRI anatomic acquisition protocols<sup>a</sup>** |  |  |  |  |  |  |
| --- | --- | --- | --- | --- | --- | --- |
|  | **3D T2 FLAIR** |  |  | **3D T1 MPRAGE** |  |  |
|  | **Siemens Skyra** | **Siemens Prisma** | **Philips Achieva** | **Siemens Skyra** | **Siemens Prisma** | **Philips Achieva** |
| Operation system version | syngo VE11C | syngo VE11C | R5.6.1 | syngo VE11C | syngo VE11C | R5.6.1 |
| Head Coil | 32 channel | 32 channel | 32 channel | 32 channel | 32 channel | 32 channel |
| Acceleration factor for parallel imaging | 2 | 2 | 2.6 | 2 | 2 | 2 |
| Orientation | Sagittal | Sagittal | Sagittal | Sagittal | Sagittal | Sagittal |
| FOV (cm) | 25.6 x 25.6 | 25.6 x 25.6 | 25.6 x 25.6 | 25.6 x 25.6 | 25.6 x 25.6 | 25.6 x 25.6 |
| Matrix Size | 256 x 256 | 256 x 256 | 256 x 256 | 256 x 256 | 256 x 256 | 256 x 240 |
| No. of Sections | 176 | 176 | 176 | 176 | 176 | 176 |
| TR (ms) | 4800 | 4800 | 4800 | 1900 | 1900 | 2500 |
| TE (ms) | 352 | 352 | 305 | 2.52 | 2.52 | 3.14 |
| TI (ms) | 1800 | 1800 | 1650 | 900 | 900 | 900 |
| Flip Angle | 120 | 120 | 90 | 9 | 9 | 9 |
| Voxel Size (mm) | 1.0 × 1.0 × 1.0 | 1.0 × 1.0 × 1.0 | 1.0 × 0.89 × 0.89 | 1.0 × 1.0 × 1.0 | 1.0 × 1.0 × 1.0 | 1.0 × 1.0 × 1.0 |
| Scan Time (min:s) | 6:57 | 6:57 | 6:33 | 4:15 | 4:15 | 5:09 |
| No. of Signal Averages | 1 | 1 | 1 | 1 | 1 | 1 |
| <sup>a</sup> The Brigham and Women's Hospital and the National Institute of Health used a Siemens Skyra scanner, the University of Pennsylvania used a Siemens Prisma scanner, and the Johns Hopkins University used a Philips Achieva Scanner. |  |  |  |  |  |  |

The unexpected activation of a single spine coil receive element due to head placement in the scanner during several imaging sessions at Penn and Brigham resulted in shading artifacts through the caudal areas of the images, as shown in **Figure 2**.

**Figure 2.**
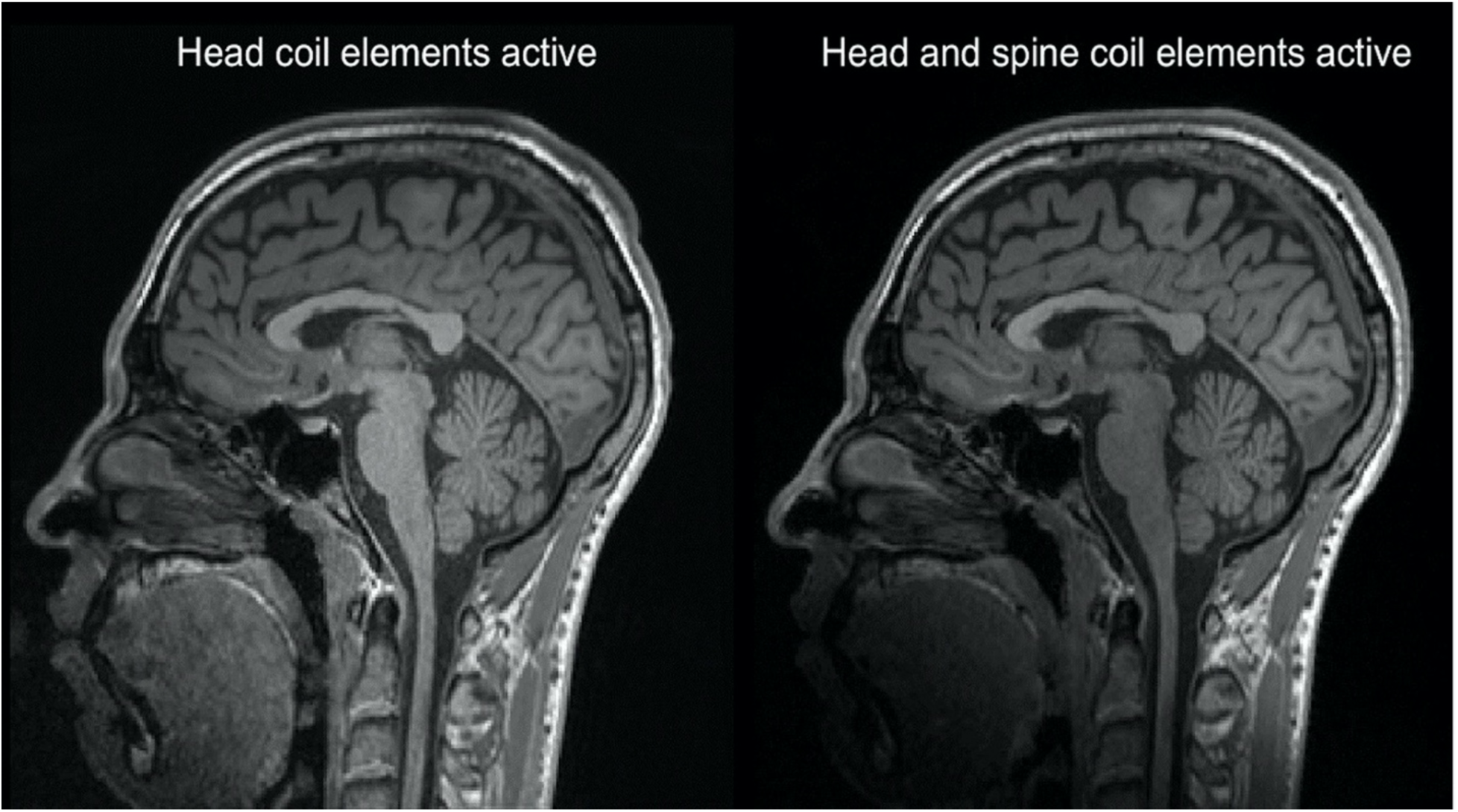
Comparison of a pair of scan-rescan images acquired during the same session on the same scanner in which head positioning resulted in a spine receive coil activation (right) compared to not (left). Note excess shading apparent in the inferior anterior region of the image resulting from spine coil activation.

### Manual Lesion Segmentation

Images were de-identified and manually assessed for identification of individual lesions and quantification/contouring of each lesion to derive total cerebral T1-hypointense lesion volume (T1LV) and T2LV from the native 3D T1 and FLAIR images, respectively. This was performed by one experienced observer (S.T.); any challenging cases were verified by a senior lab director (R.B.). For T2LV, all lesions on the FLAIR images were identified. For T1LV, lesions showing both hypointensity on T1-weighted images and at least partial hyperintensity on FLAIR images were marked. A semiautomated edge-finding tool in Jim (Version 7.0; http://www.xinapse.com/home.php) was then employed for delineation and volume estimation. De-identified images were pooled for the entire study, analyzed in one batch, and were presented in random order to reduce scan-to-scan memory effects. Estimated volumes obtained using Jim are shown in **Figure 3**.

**Figure 3.**
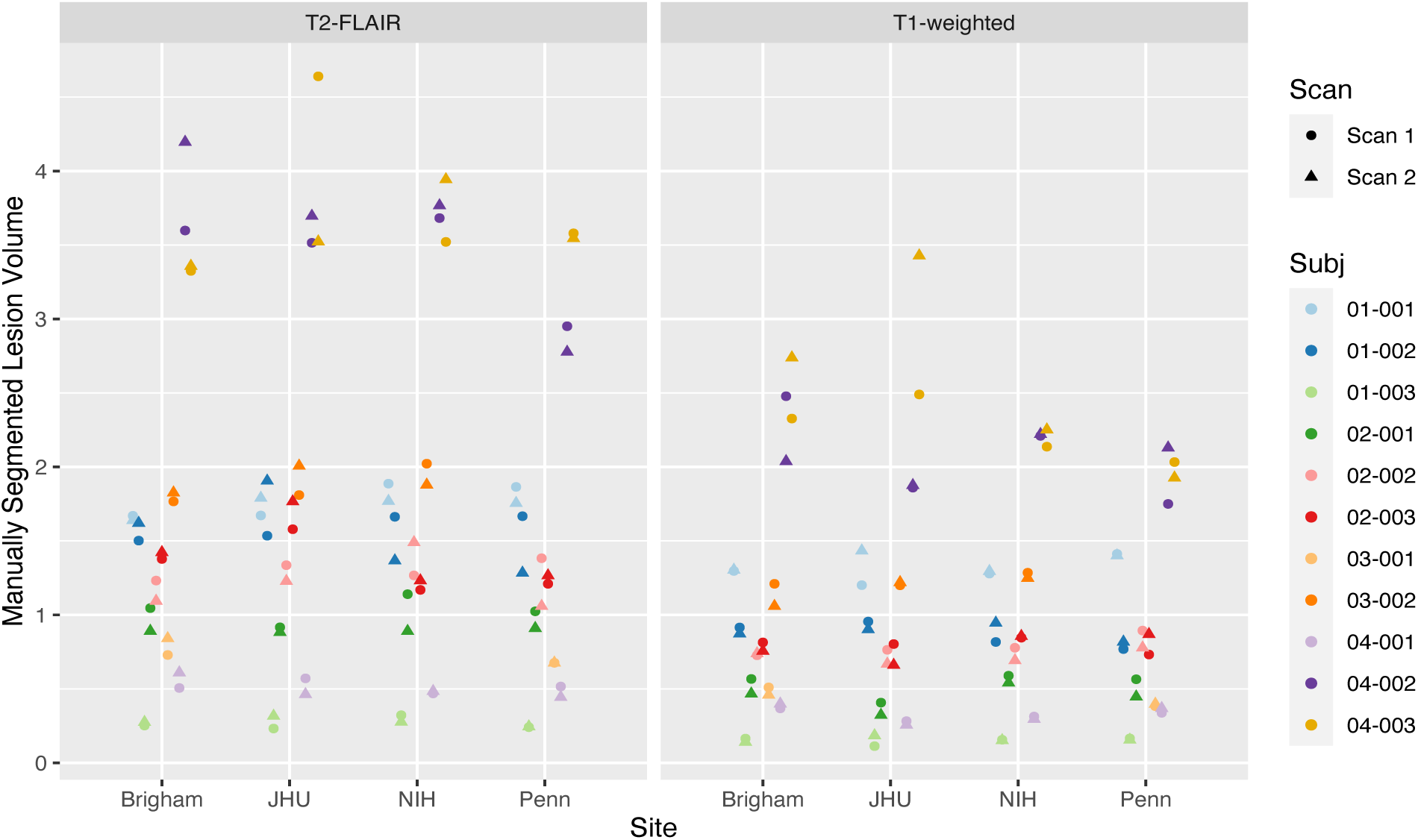
Manually measured T1-weighted and T2-FLAIR lesion volumes for scan-rescan pairs of images from 11 subjects at each of the 4 NAIMS sites. Scan 1 volumes are indicated by circles, and scan 2 volumes are shown with triangles. Each subject is represented by a different color, and points are offset from one another to aid in visualization.

### Pre-processing

Prior to automated segmentation, images underwent bias correction via nonuniform intensity normalization (N4ITK),^21^ and FLAIR images were rigidly aligned to the subject’s own corresponding T1-weighted image within a given scan session. Brain extraction was performed using Multi-Atlas Skull Stripping (MASS),^22^ and intensity normalization was performed using WhiteStripe prior to automated lesion segmentation.^23^

### Automated Volumetry Methods

Fully automated WM lesion segmentation was performed using the Method for Inter-Modal Segmentation Analysis (MIMoSA), a logistic regression-based WM lesion segmentation method that incorporates mean structure and local covariance structures across imaging modalities obtained by intermodal coupling.^24^ Normal appearing white matter and gray matter volumes were estimated using Joint Label Fusion (JLF), a multi-atlas segmentation method that utilizes weighted voting based on voxel-level joint probability of a segmentation error occurring in pairs of atlases in order to minimizes total labelling error.^25^ Whole brain volume was estimated using FMRIB’s Automated Segmentation Tool (FAST).^26^ Estimated thalamic volumes were obtained using JLF and FMRIB’s Integrated Registration and Segmentation Tool (FIRST).^27^

### Statistical Analysis

All statistical analyses and visualization were performed in R (version 4.1.1) (http://www.r-project.org/). Random effects modeling was performed using the lme4 package (version 1.1.27.1).^28^ Models utilizing a classical random intercept structure, which included a random effect term that nested site within subject (to account for the interaction between site and subject as well as subject-level clustering for each of our brain structures and volumetry methods), were used to assess site biases in estimated volumes. Validity of the random effects models was assessed visually using normal Q-Q plots of the residuals and random effects created using the stats package (version 4.1.1), residual versus fitted plots, and density plots of the random effects for each model using the ggplot2 package (version 3.3.5). The Shapiro-Wilk test was used to assess normality of the residuals for each model using the stats package.

Linear regression was performed using the stats package to model the relationship between site and estimated volumes within each subject for all brain structures and volumetry methods, and the proportion of variation explained by site was computed to assess the effect of site on estimated volumes within each subject from these models.

To complement the parametric modeling approach, mean absolute difference (MAD) across all subjects was computed within and across sites. This was done by computing the absolute differences between all possible pairs of intra-scanner measures and inter-scanner measures within each subject. These differences were then averaged across all subjects to obtain a measure of the mean absolute inter-scanner difference and mean absolute intra-scanner difference followed by a calculation of a ratio of these two measures. Permutation testing was performed to assess whether the MADs of inter- and- intra-scanner measures were significantly different, using the ratio of MADs across inter-scanner measures and intra-scanner measures as our test statistic. Under the null hypothesis, we would expect a ratio of 1, indicating that average inter-scanner and average intra-scanner differences are equal. Ten-thousand permutations were performed, which involved shuffling site labels across all measures within each subject, and the ratio of average absolute inter- and intra-scanner differences was computed after each permutation.

Biases in estimated volumes associated with active coil elements were assessed by computing the median absolute difference between estimated volumes for a) within-subject pairs of images that were acquired using the same active coil elements (namely, either both with or without activation of a spine coil element) and b) within-subject pairs of images that were acquired using different active coil elements was computed for all ROIs and methods. These were then averaged across all subjects. Permutation testing was performed to assess whether the median absolute differences from measures acquired using similar active coil and different active coil elements were significantly different, using the ratio of median absolute difference across different coil measures and median absolute difference across same coil measures as our test statistic. 10,000 permutations were performed by shuffling coil labels within each site (Penn and Brigham), and the ratio described above was computed after each permutation.

## Results

Visual inspection of the residuals and effects from the random effect models indicated that most of the models violated the assumptions of normal distribution and homogeneity of the residuals which is exemplified in **Figure 4** as well as normal distribution of the random effects in some models. This led to marked underestimation of the proportion of variance explained by site.

**Figure 4.**
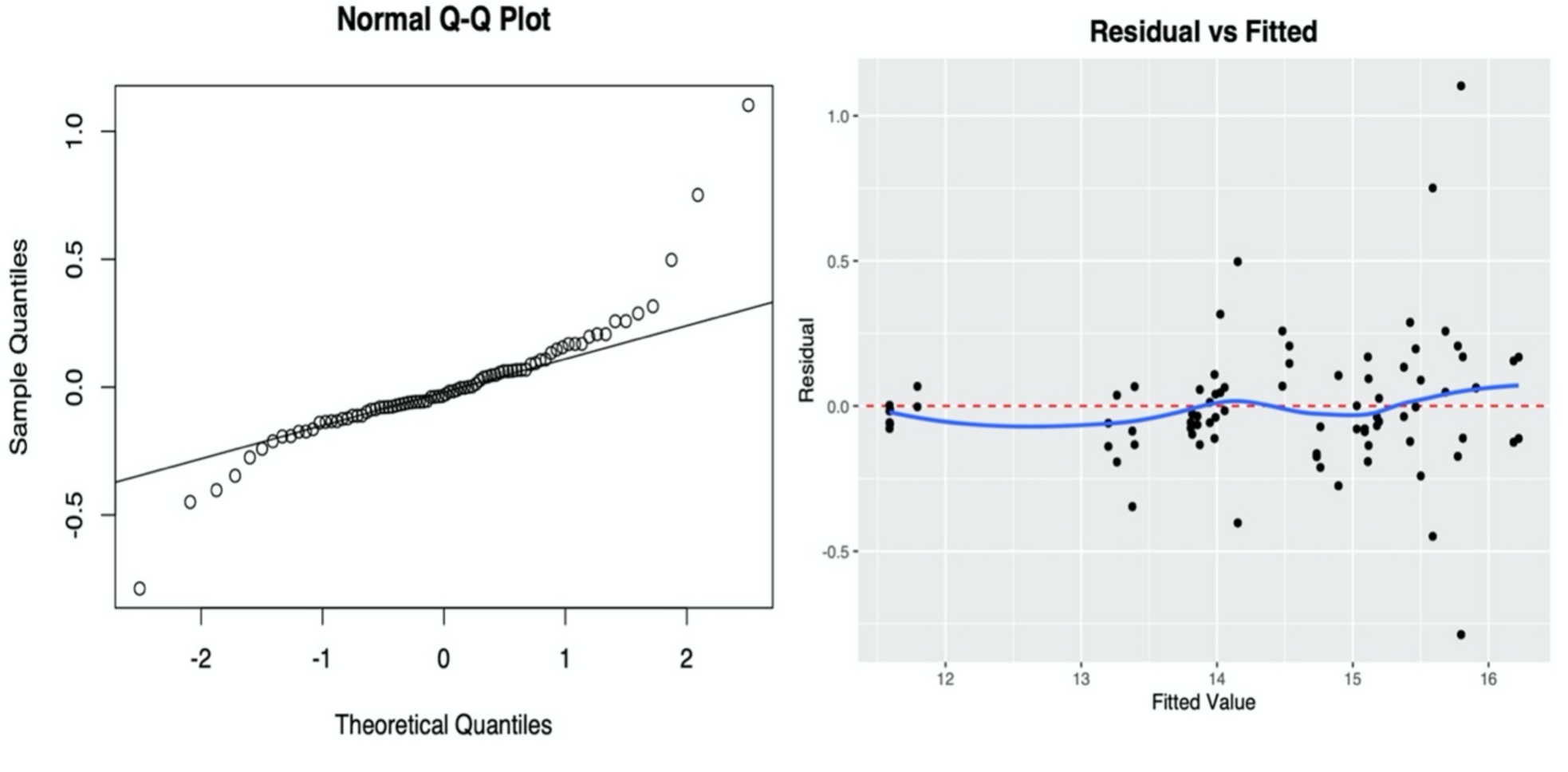
Normal Q-Q and residual versus fitted plots for the random effect model which includes thalamic volumes obtained using JLF from distortion corrected images acquired at all 4 sites as the response variable and a random effect term which nested site within subject. The heavy tailed distribution of the Normal Q-Q plot indicates non-normal distribution of the residuals from the random effect model. As the fitted values increase in the residual vs. fitted plot, variance of the residuals also increases indicating heteroskedasticity of the residuals.

The spread of the proportion of variation explained by site from linear models for each brain structure and volumetry methods across all subjects is shown in **Figure 5**. Site explained > 50% of the variation in most subjects across most automated methods. More than 75% of the variation was explained by site in most subjects for most methods and brain structures where data from all scanner manufacturers was included in the models, indicating notable differences across scanner manufacturers.

**Figure 5.**
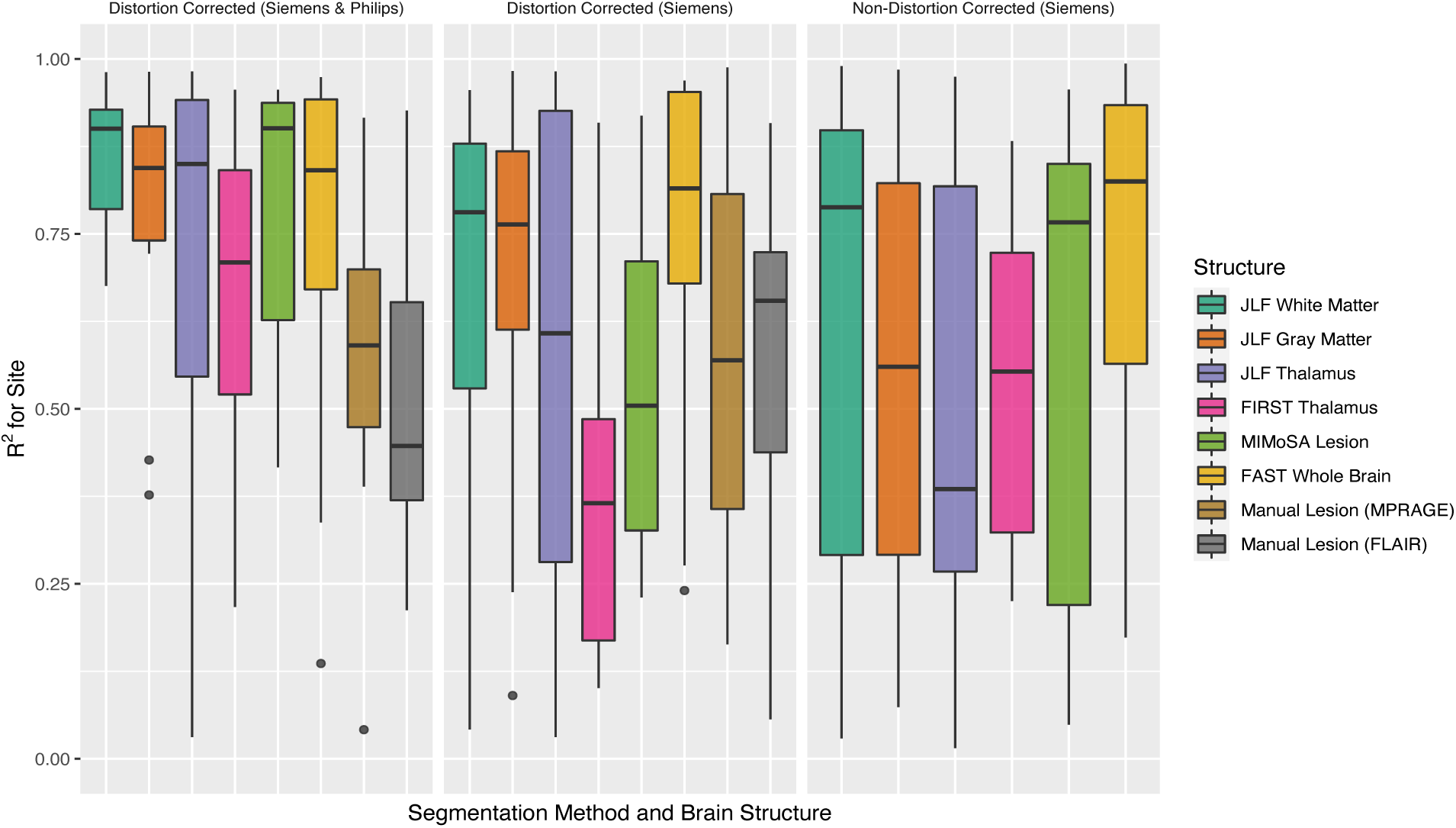
Proportion of variation explained by site in brain volumes extracted via various methods represented by different colors. Each panel indicates settings which either included or excluded data from the Philips scanner as well as the distortion correction settings. Each boxplot depicts the spread of R^2^ values for site across all subjects.

Ratios of average absolute inter- and intra-scanner differences are shown in **Figure 6**. Mean absolute differences (MAD) across pairs of inter-scanner measures were greater compared to intra-scanner measures for most volumetry methods and brain structures, with the largest ratios of average differences ranging from approximately 1.5 to 2.5 and occurring in subgroups that included data acquired from both scanner manufacturers. This indicates larger inter-scanner differences in volumes acquired on both Siemens and Phillips scanners compared with those acquired on only Siemens scanners for most of the automated volumetry methods.

**Figure 6.**
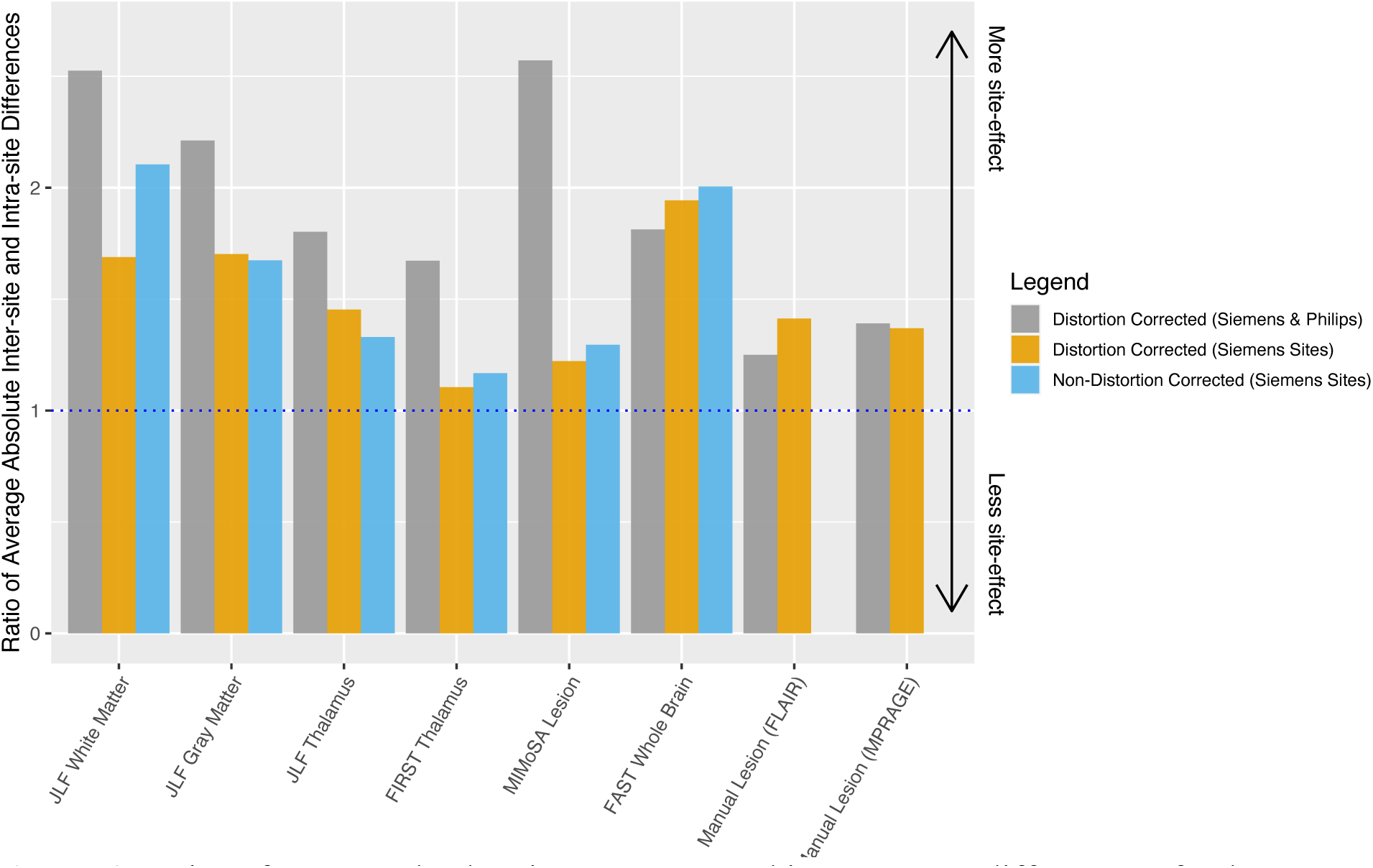
Ratios of average absolute inter-scanner and intra-scanner differences of volumes extracted via various methods (horizontal axis). Colors indicate different settings which either included or excluded data from the Philips scanner, as well as the distortion correction setting. The blue dashed line represents where inter-scanner and intra-scanner differences are equal; inter-scanner differences were larger than intra-scanner differences most measures.

Average absolute difference between all possible inter-subject pairs of images was computed, and ratios of average absolute inter-subject and inter-scanner differences are shown in **Figure 7**. Average inter-subject differences were larger than average inter-scanner differences, ranging from approximately 1.5 to 10 times larger across all volumetry methods and brain structures, indicating that differences between subjects were of greater magnitude than inter-scanner differences.

**Figure 7.**
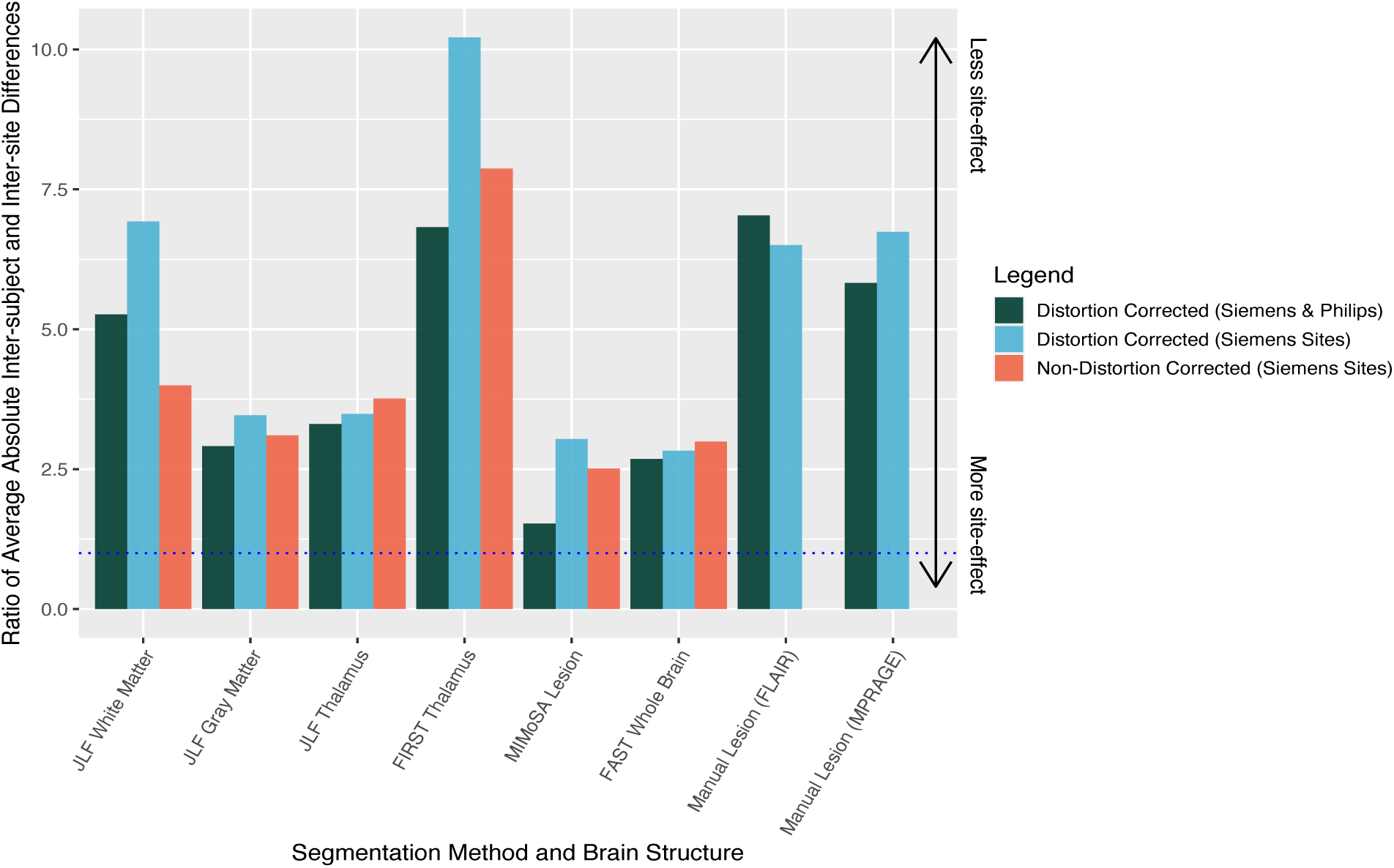
Ratios of average absolute inter-subject and inter-scanner differences of volumes extracted via various methods (horizontal axis). Colors indicate different settings which either included or excluded data from the Philips scanner, as well as the distortion correction setting. The blue dashed line represents where inter-subject and inter-scanner differences are equal; inter-subject differences were larger than inter-scanner differences for all measures.

Permutation testing was performed to assess whether the MADs of inter- and- intra- scanner measures were significantly different, using the ratio of MADs across inter-scanner measures and intra-scanner measures as our test statistic. Under the null hypothesis, we would expect a ratio of 1, indicating that average inter-scanner and average intra-scanner differences are equal. 10,000 permutations were performed, involving the shuffling of site labels within each person, and the ratio of average absolute inter- and intra-scanner differences was computed after each permutation. The resulting negative log (base 10) *p-*values from these permutation tests are shown in **Figure 8**. Significant *p*-values were observed for many of the automated volumetry methods and brain structures both before and after multiple comparison correction. Most of these corresponded to tests that included data from both Siemens and Philips sites, indicating that significant differences in volumes were most prevalent across different scanner manufacturers.

**Figure 8.**
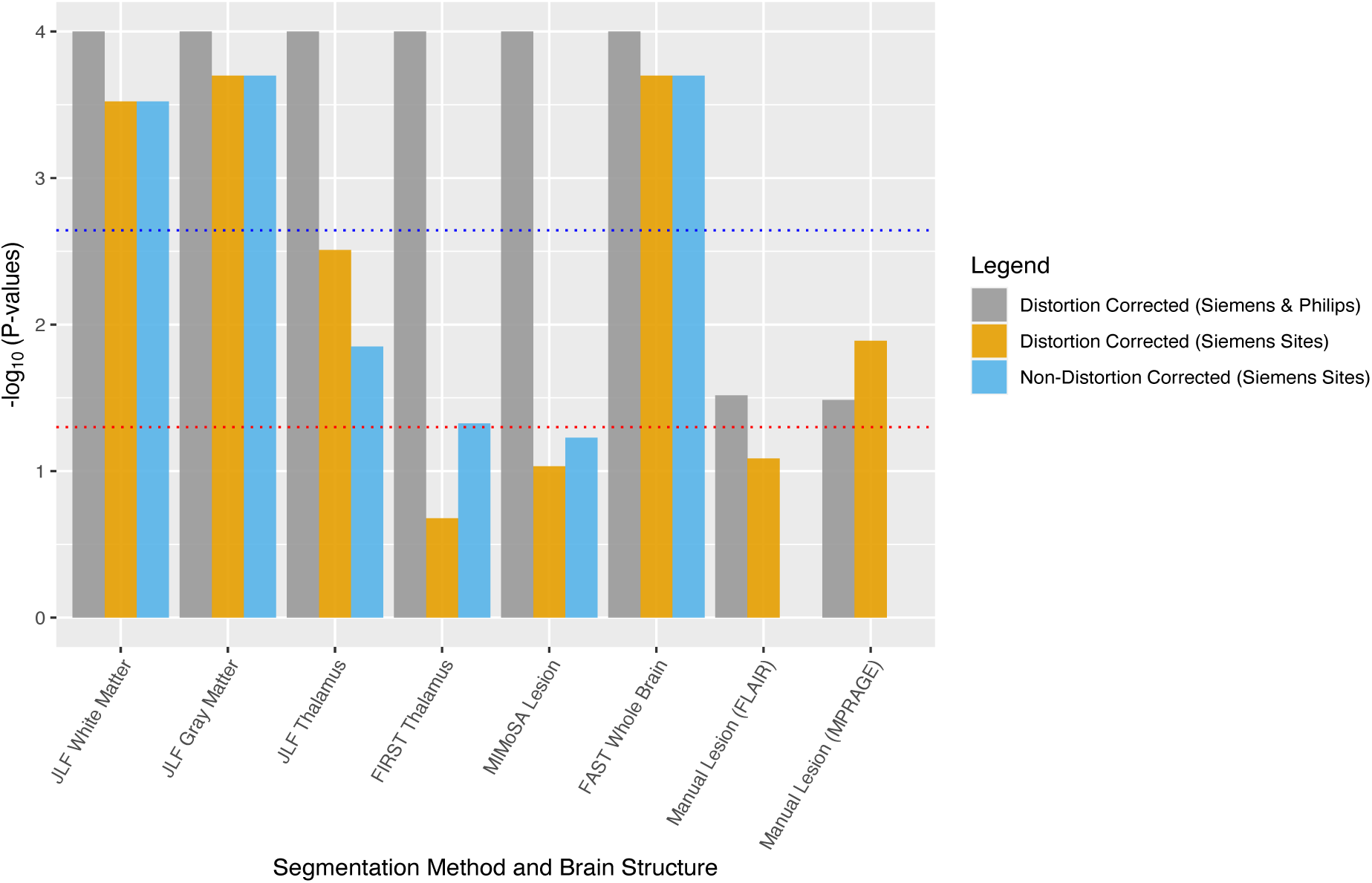
Negative log (base 10) *P* values from permutation tests assessing the association between site and brain volume extracted via various methods (horizontal axis). Colors indicate different settings which either included or excluded data from the Philips scanner, as well as the distortion correction setting. The red dashed line represents the unadjusted significance threshold of 0.05, and the blue dashed line represents the significance threshold obtained from the Bonferroni correction method. Most measurements demonstrated significant site effects, with many settings surviving even conservative Bonferroni correction.

To assess differences in estimated volumes associated with active coil elements during imaging, permutation testing was performed using the ratio of median absolute differences across all pairs of within subject measures from images acquired using different active coil elements and those acquired using the same active coil elements and the resulting negative log (base 10) *p-*values from these permutation tests are shown in the Online Supplemental Data **On- line Figure 5**. Distortion and non-distortion-corrected imaging indicated multiple comparison-adjusted significant differences in JLF-measured thalamic volumes across receive coil modes (p < 0.01), potentially due the proximity of the thalamus to the receive coil shading artifact and JLF is more sensitive to global registration differences.

## Discussion

The aim of this study was to quantify differences in brain segmentation volumes from patients with MS across different scanners and sites to further understand how site biases manifest in the presence of varying MS pathology. The use of a standardized high-resolution protocol at 3T across all sites was crucial in minimizing the effect of scanning parameters on variation in estimated volumes. Critically, all study participants were clinically stable at the time of inclusion and radiologically stable throughout the imaging period, and imaging was completed within approximately 3 months of each person’s initial imaging session to minimize the effect of biological variation on estimated volumes. Moreover, as previous studies have demonstrated an association between brain volumes and time of day and hydration status,^29–31^ imaging sessions were scheduled within a consistent time of day across all visits for each subject in order to control for this potential source of variation.

Biases in estimated volumes were assessed across several ROIs of key interest in MS using an array of automated methods to improve generalizability of our results. Despite these considerations, biases in brain volumes associated with site persisted and were found to be most notable across sites where different scanner vendors were used.

Further study is necessary to develop an appropriate parametric model to represent these site biases, since random and mixed effects modelling failed due to violations of classical statistical assumptions. Non-normality of the residuals and random effects from models that included a nested random effect to account for the interaction between site and subject led to underestimation of the variance in estimated volumes explained by site for most ROIs/methods considered in this analysis.

Head placement in the scanner resulted in the unanticipated activation of a spine receive coil element in 2 Siemens scanners, which produced a shading artifact unique to this occurrence. Permutation testing revealed that median absolute difference across pairs of estimated volumes measures from images where active coil elements differed during imaging was significantly more than that of images acquired using similar active coil elements in the case of thalamic volumes. These findings warrant further investigation of the effect of active receive coil elements on volumetry and suggest that consideration of active receive coils employed, beyond the physical receive array device, and their settings in scanning protocols could reduce bias both within and across scanners.

This analysis involved a limited number of participants and range of scanners, and we obtained research-quality images from a small number of large academic hospitals, which does not reflect the level of variation we might expect to see across additional participants, scanners, and community hospitals and independent radiology practices. Further analysis involving a larger number of participants, scanners, vendors and versions, sites, and longitudinal imaging follow-up is warranted to assess generalizability of our results and to understand how site effect in our analysis relates to site effect in longitudinal clinical follow-up.

## Conclusion

Imaging 11 people with stable MS within 3 months of their initial study visit at 4 different NAIMS sites allowed us to assess inter- and intra-scanner differences in brain volumetric measurements. Significant technical variation in estimated volumes due to site were present across most ROIs and automated methods despite careful protocol harmonization. Average inter-scanner differences were largest relative to intra-scanner differences in subgroups that included data from both Philips and Siemens scanners, indicating higher variability across scanner manufacturer compared to variability in estimated volumes from images acquired using the same scanner manufacturer. Automatic activation of a receive coil element in the spine coil on Siemens scanners contributed to increased significant biases in volumetric measurements.

Our results highlight the persistence of inter-scanner variation even when using a harmonized protocol and stress the need to account for inter-scanner variation in clinical and research settings as they have the potential to confound study results. In cases where automated volumetry is used in clinical decision making, the effects such variation should be considered when making treatment decisions. Further study is warranted to improve our understanding of site effect in people with MS and to develop methods to mitigate these site effects.

## Supporting information

Supplemental On-line Figures

## Acknowledgements

We acknowledge the contribution of the NINDS Neuroimmunology Clinic and the NIMH Functional Magnetic Resonance Facility

