## Supplemental On-line Figures for "Inter-scanner brain MRI volumetric biases persist even in a harmonized multi-subject study of multiple sclerosis"

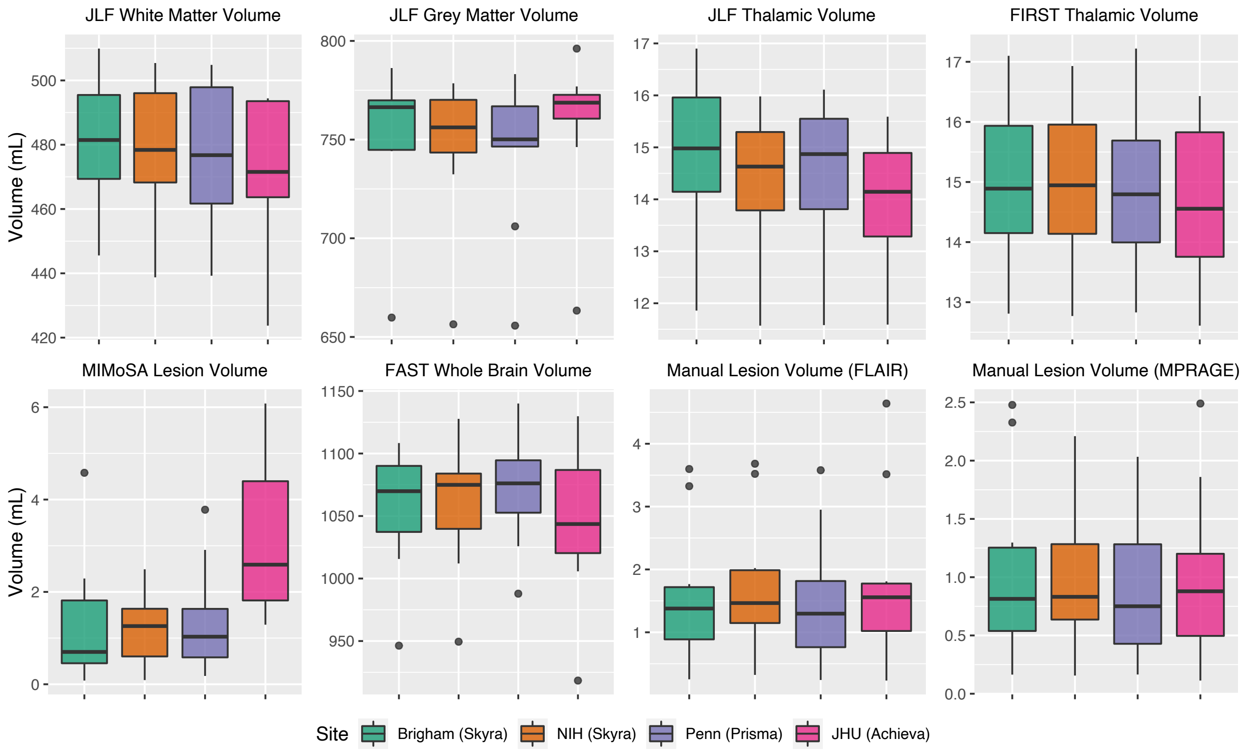


**On-line Figure 1.** Comparison of estimated volumes from distortion-corrected scan 1 images for all 11 subjects across all 4 sites depicting inter-subject variability within each site as well as inter-scanner variability for all estimated volumes..

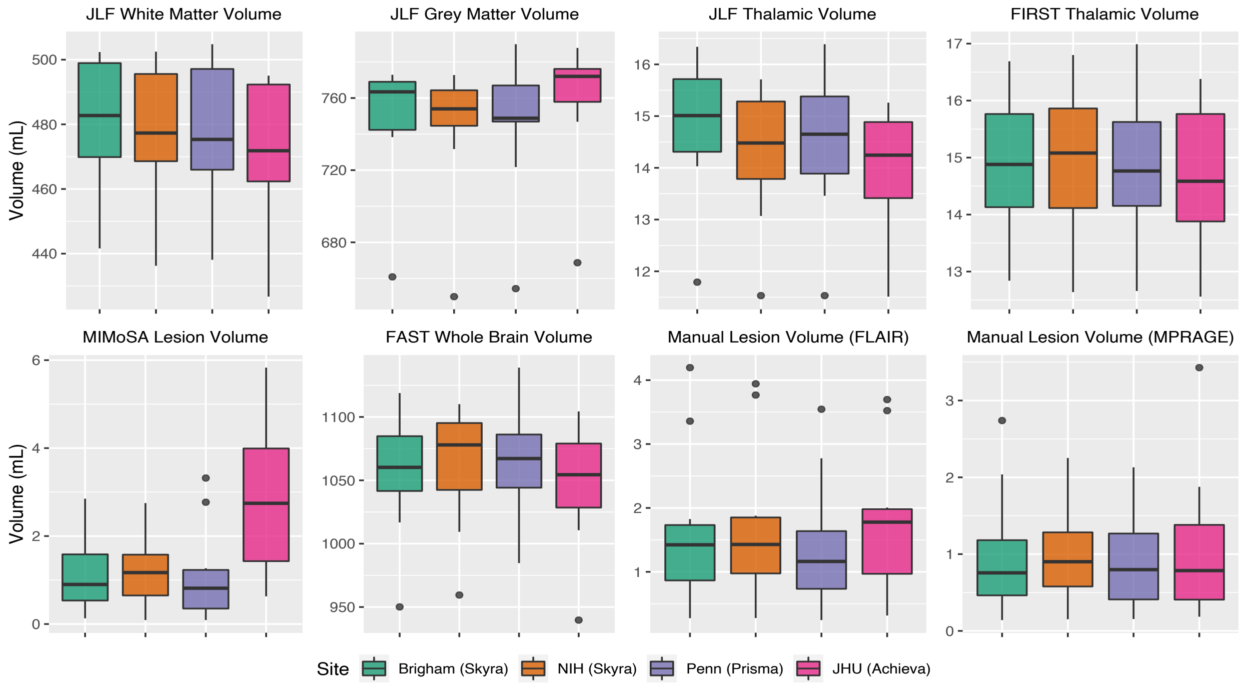


**On-line Figure 2**. Comparison of estimated volumes from distortion-corrected scan 2 images for all 11 subjects across all 4 sites depicting inter-subject variability within each site as well as inter-scanner variability for all estimated volumes.

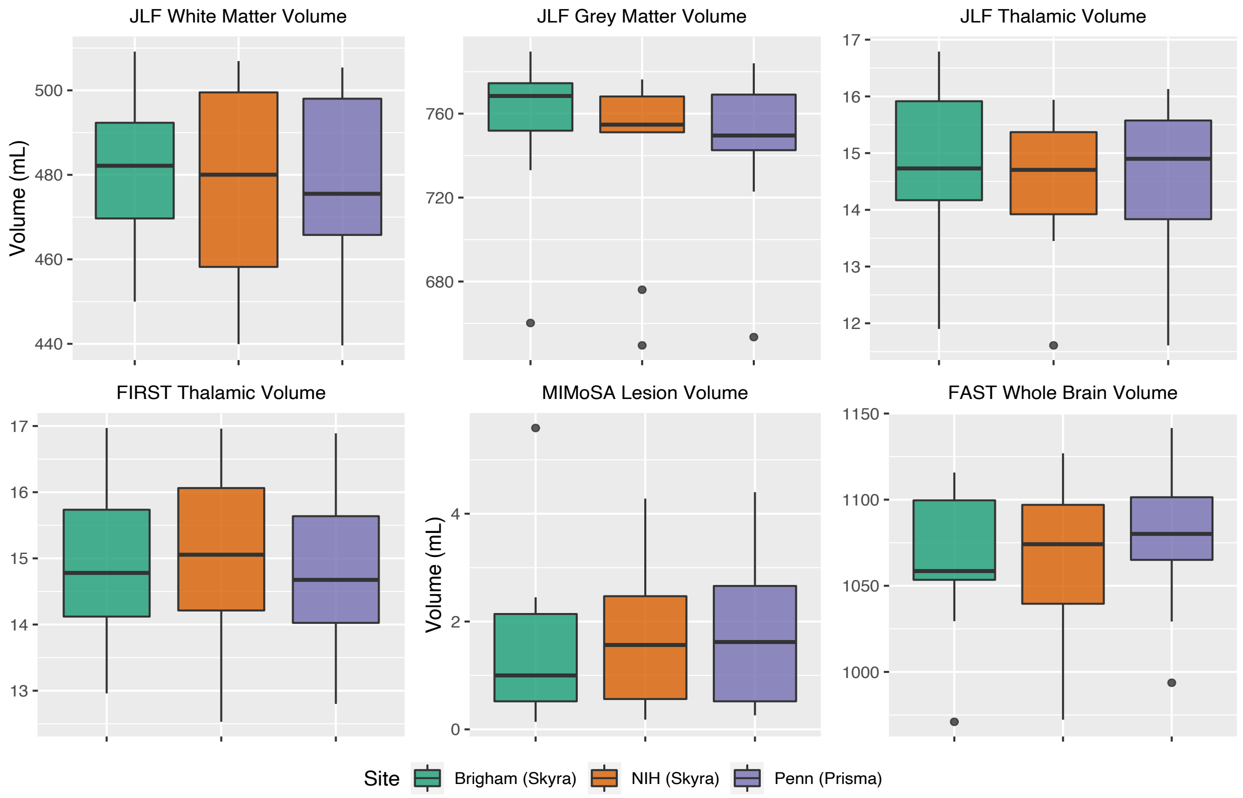


**On-line Figure 3.** Comparison of estimated volumes from non-distortion corrected scan 1 images for all 11 subjects across the 3 sites that reconstructed non-distortion corrected images depicting inter-subject variability within each site as well as inter-scanner variability for all estimated volumes.

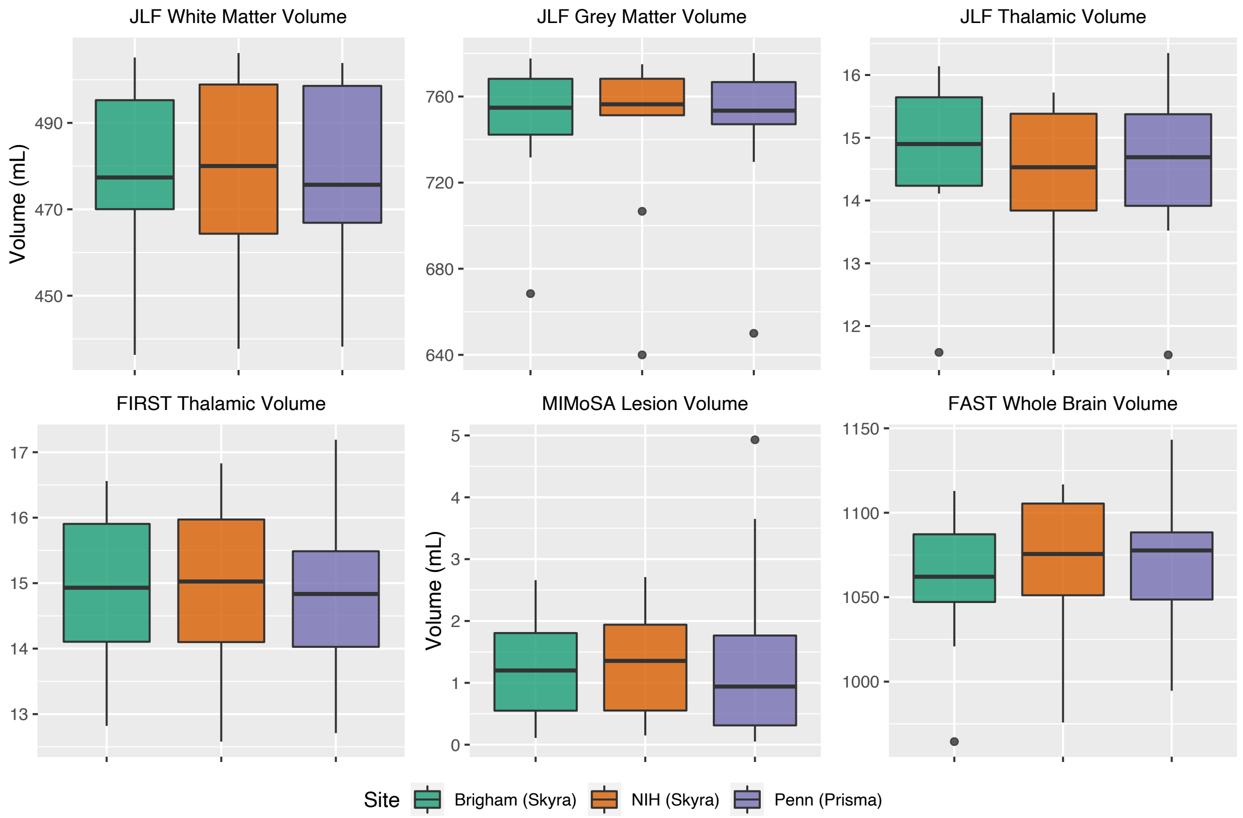


**On-line Figure 4.** Comparison of estimated volumes from non-distortion corrected scan 2 images for all 11 subjects across the 3 sites that reconstructed non-distortion corrected images depicting inter-subject variability within each site as well as inter-scanner variability for all estimated volumes.

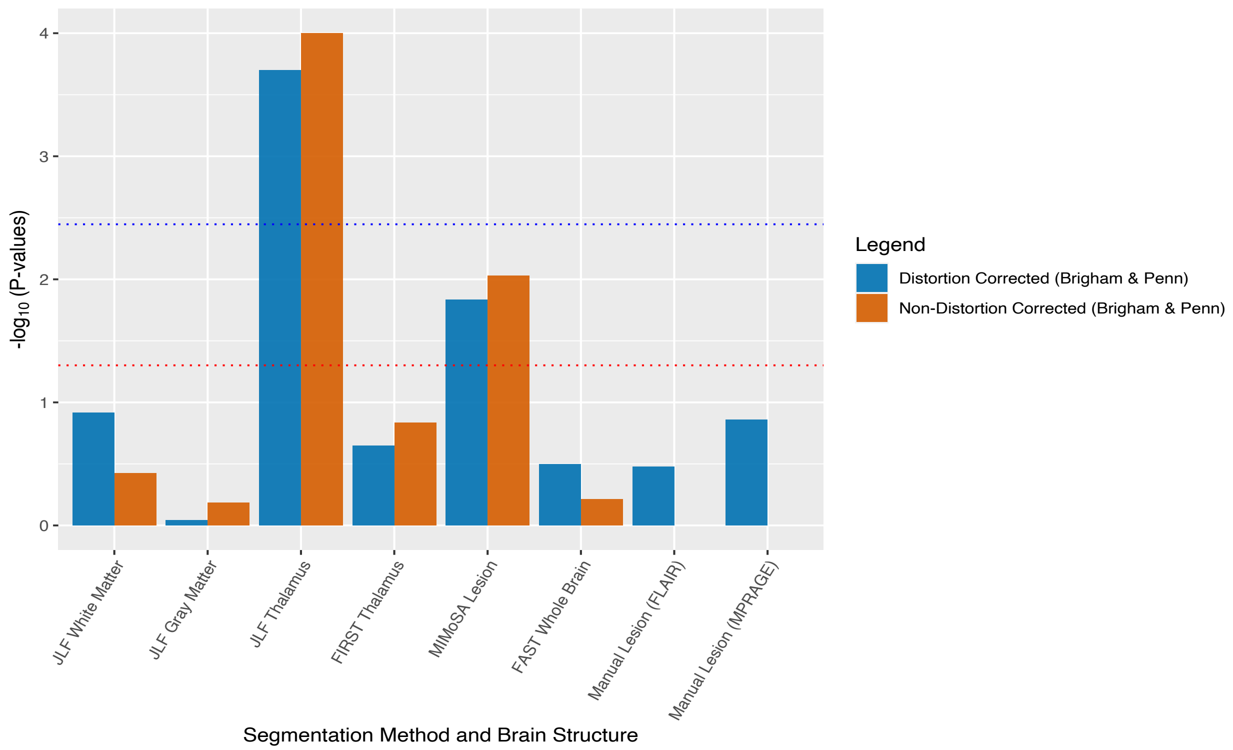


**On-line Figure 5.** Negative log (base 10) *P* values from permutation tests assessing the association between active coil elements during imaging and brain volumes extracted via various methods (horizontal axis). Colors indicate the distortion correction setting. The red dashed line represents the unadjusted significance threshold of 0.05, and the blue dashed line represents the significance threshold obtained from the Bonferroni correction method. Thalamic measurements using JLF and MIMoSA lesion volumes demonstrated significant effects, with JLF thalamus results surviving Bonferroni correction.
